# Metformin protects trabecular meshwork against oxidative injury via activating integrin/ROCK signals

**DOI:** 10.1101/2022.06.30.498232

**Authors:** Lijuan Xu, Xinyao Zhang, Yin Zhao, Yang Cao, Yuanbo Liang

**Affiliations:** National Clinical Research Center for Ocular Diseases; The Eye Hospital of Wenzhou Medical University; Glaucoma Research Institute of Wenzhou Medical University, Zhejiang, China

**Keywords:** primary open-angle glaucoma, trabecular meshwork, cytoskeleton remodelling, metformin

## Abstract

**Background:** This study aimed to investigate the protective effect of metformin on the trabecular meshwork (TM) and explore its molecular mechanisms *in vivo* and *in vitro*.

**Methods:** Ocular hypertension (OHT) mouse models were induced with dexamethasone (DEX) and further treated with metformin to determine its IOP lowering effect. Cultured human TM cells (HTMC) were pre-stimulated with tert-butyl hydroperoxide (tBHP) to induce oxidative damage and then supplemented with metformin for another 24 h. The expression of fibrotic markers and integrin/ROCK signals, including α-SMA, TGF-β, fibronectin, F-actin, integrin beta 1, Rho-associated kinase (ROCK)1/2, AMP-activated protein kinase (AMPK), myosin light chain 1 (MLC 1), and F-actin were determined by western blotting (WB) and immunofluorescence (IF). Reactive oxygen species (ROS) content was analysed using flow cytometry (FCM).

**Results:** Administration of metformin reduced the elevated IOP and alleviated the fibrotic activity of aqueous humour outflow in OHT models. Additionally, metformin rearranged the disordered cytoskeleton in the TM both *in vivo* and *in vitro*. Furthermore, metformin significantly inhibited ROS production and activated integrin/ROCK signalling induced by tBHP in HTMC.

**Conclusion:** Metformin reduced the elevated IOP in steroid-induced OHT mouse models and exerted its protective effects against oxidative injury by regulating cytoskeleton remodelling through the integrin/ROCK pathway. This study provides new insights into metformin use and preclinical evidence for the potential treatment of primary open-angle glaucoma.

## Introduction

Intraocular pressure (IOP) elevation, predominantly resulting from increased resistance to aqueous humour outflow (AHO), is a major risk factor for primary open-angle glaucoma (POAG) deterioration (*Casson, et al.,2012*; *Wu, et al.,2020*). The only proven method is IOP lowering (*Richter and Coleman,2016*). According to Bill and his colleagues (*Bill and Hellsing,1965*; *Bill and Svedbergh,1972*), up to 80% aqueous humour is drained via conventional trabecular meshwork (TM) pathway; however, the available anti-glaucoma medications mostly act on sites other than TM and have limited efficiency. Therefore, polypharmacy has become increasingly prevalent, coupled with an increasing economic burden on society and patients (*Wu, et al.,2020*).

IOP elevation is associated with TM stiffness (*Alvarado, et al.,1984*; *Heijl, et al.,2002*; *Johnstone, et al.,2021*). Theoretically, TM-targeting drugs are potentially effective in lowering IOP caused by diseased TM. Integrin and Rho-associated protein kinase (ROCK) play pivotal roles in cytoskeleton formation and maintenance (*Tan, et al.,2020*; *Yemanyi, et al.,2020*) and ROCK inhibitor (ROCKi) can decrease actomyosin contraction and actin crosslinking (*Liu, et al.,2021*). ROCKi is the only drug that directly targets conventional outflow function (*Aga, et al.,2008*; *Rao, et al.,2001*; *Ren, et al.,2016*). It alters the architecture of AHO and expands the juxtacanalicular connective tissue region. Currently, the clinically available ROCKi include Y-27632 and ripasudil. However, the costs of Y-27632 and ripasudil are as high as 300 $ (2.5 mL) and 209 $ (5 mL), respectively. Moreover, they have not yet been approved for use in China. Considering that China is a developing country with the largest population in the world, it is urgent to explore novel TM-targeting drugs with characteristics of efficiency and lower prices, which are appropriate for our nation.

Metformin (MET), an oral biguanide insulin-sensitising drug, is the most widely used treatment for type 2 diabetes mellitus (DM) (*Foretz, et al.,2014*). It is a multifunctional drug (*Rangarajan, et al.,2018*; *Zhao, et al.,2021*). Recent studies by Lin et al. (*Lin, et al.,2015*) and Maleskic et al. (*Maleskic, et al.,2017*) found that metformin reduced the risk of open-angle glaucoma (OAG) in patients with DM, and this effect persisted even after controlling for glycated haemoglobin. However, this effect was not observed with other hypoglycaemic medications (insulin, sulfonylureas, thiazolidinediones, and meglitinides), suggesting that the protective effect of MET on glaucoma goes beyond glycaemic improvement. However, the precise mechanisms involved remain unclear.

Excessive reactive oxygen species (ROS) in TM play an important role in the disruption of cytoskeletal integrity and apoptosis (*Hu, et al.,2017*), leading to pathological alterations in AHO and subsequent IOP rise (*Babizhayev and Bunin,1989*; *Sacca, et al.,2016*). To investigate the role of metformin in damaged TM cells and ocular hypertension (OHT) mouse models, we used tert-butyl hydroperoxide (tBHP) to induce oxidative damage in TM cells (*Tang, et al.,2013*; *Wang, et al.,2021*) and topical glucocorticoids to create OHT mouse models (*Li, et al.,2021*). The results showed that metformin protected against cytoskeletal destruction in TM by enhancing the integrin/ROCK pathway and alleviated elevated IOP in steroid-induced OHT mouse models.

## Results

### Steroid-induced OHT in mouse

A steroid-induced OHT mouse model was successfully established in this study. The baseline IOP did not differ between DEX-treated and PBS vehicle eyes; however, starting at day 10, IOP was significantly elevated in DEX treated eyes (*p* < 0.05, Figure 1A) and stabilised at day 21. There were no significant differences in the initial and final levels of body weight (BW) between the two groups (Figure 1B).

**Figure 1.**
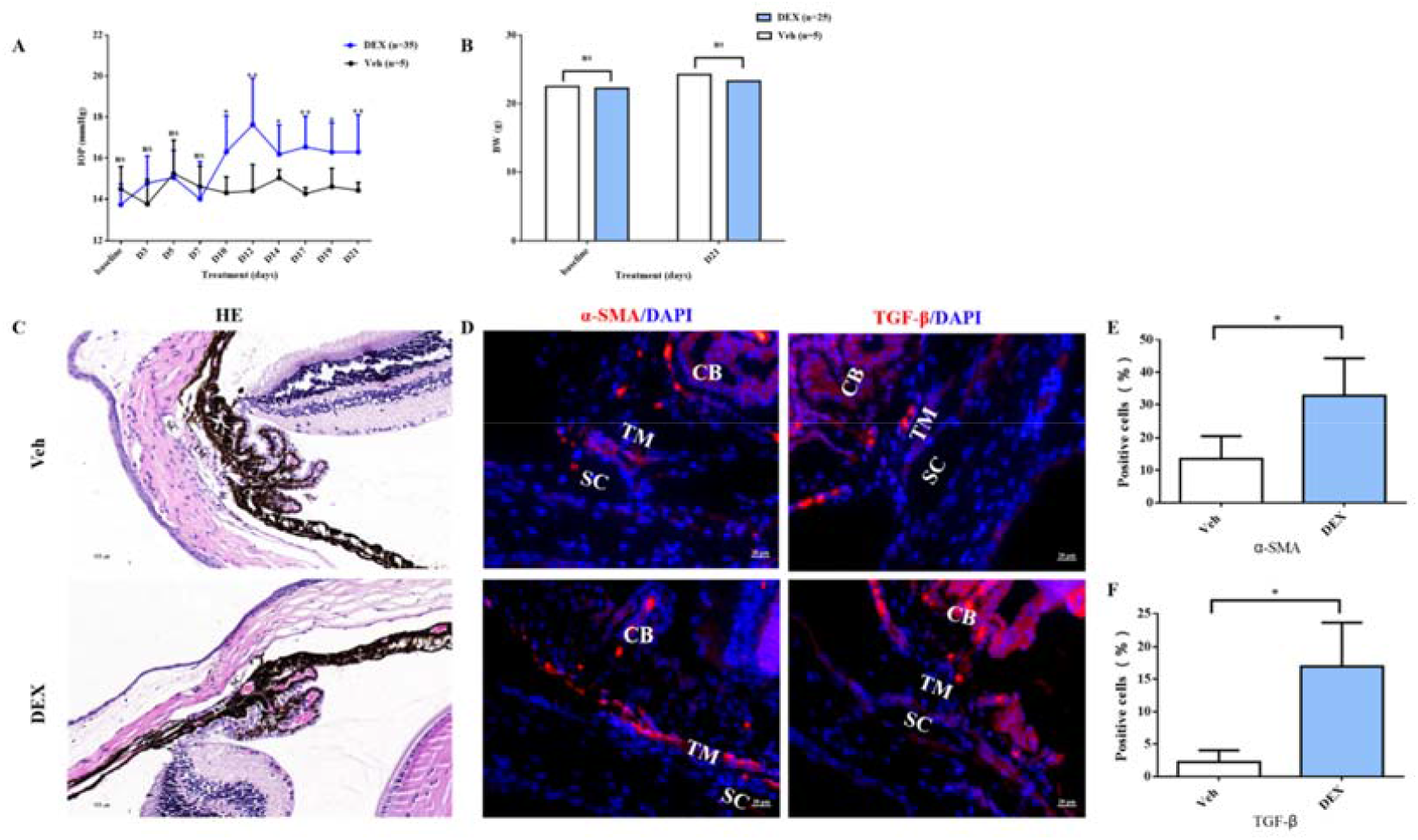
Topical ocular DEX induced OHT in mice. A. Elevated IOP in DEX-treated C57BL/6J mice was induced significantly at 3 weeks (p < 0.01). B. There was no significant change in the body weight of the DEX-treated group (*p* > 0.05). C. HE staining of OHT models. D. α-SMA and TGF-β staining in the representative OHT models. E-F. Quantification of α-SMA and TGF-β of the models. \**p* < 0.05, ns: non-significance, TM: trabecular meshwork, SC: Schlemm’s canal, CB: cililary body.

Approximately 87.5% DEX-treated eyes had elevated IOP after 21 days of steroid induction with the averaged increased magnitude of 2.73±2.21 mmHg (0.3–6 mmHg). Moreover, the fibrotic markers (α-SMA and TGF-β) were obviously overexpressed in the OHT animal models (*p* < 0.05, Figure 1D-F).

### Metformin effectively reversed steroid-induced OHT in mouse

To test the drug’s IOP lowering effect, successfully induced OHT mouse models were randomly assigned to three groups according to different treatments for an additional 21 days consecutively: PBS eye drop (group 1), 0.3% MET eye drop (group 2), and MET oral groups (group 3). Eye drops were prescribed twice daily, whereas in group 3, a certain amount of MET for each mouse was dissolved in daily drinking water.

The experimental procedure is illustrated in Figure 2A. The IOPs after 21 days of steroid induction in all three OHT groups were similar (16.67 ± 2.28, 16.62 ± 0.81, and 15.05 ± 0.88 mmHg, respectively). After five days of MET treatment, IOPs was significantly reduced in both groups 2 and 3 (*p* < 0.01, Figure 2B). In fact, metformin almost completely reversed steroid-induced OHT, returning IOP to near baseline levels on day 5 in group 3 and on day 10 in group 2, suggesting a therapeutic role of metformin in OAG. Specific IOP values are shown in Figure 2B.

**Figure 2.**
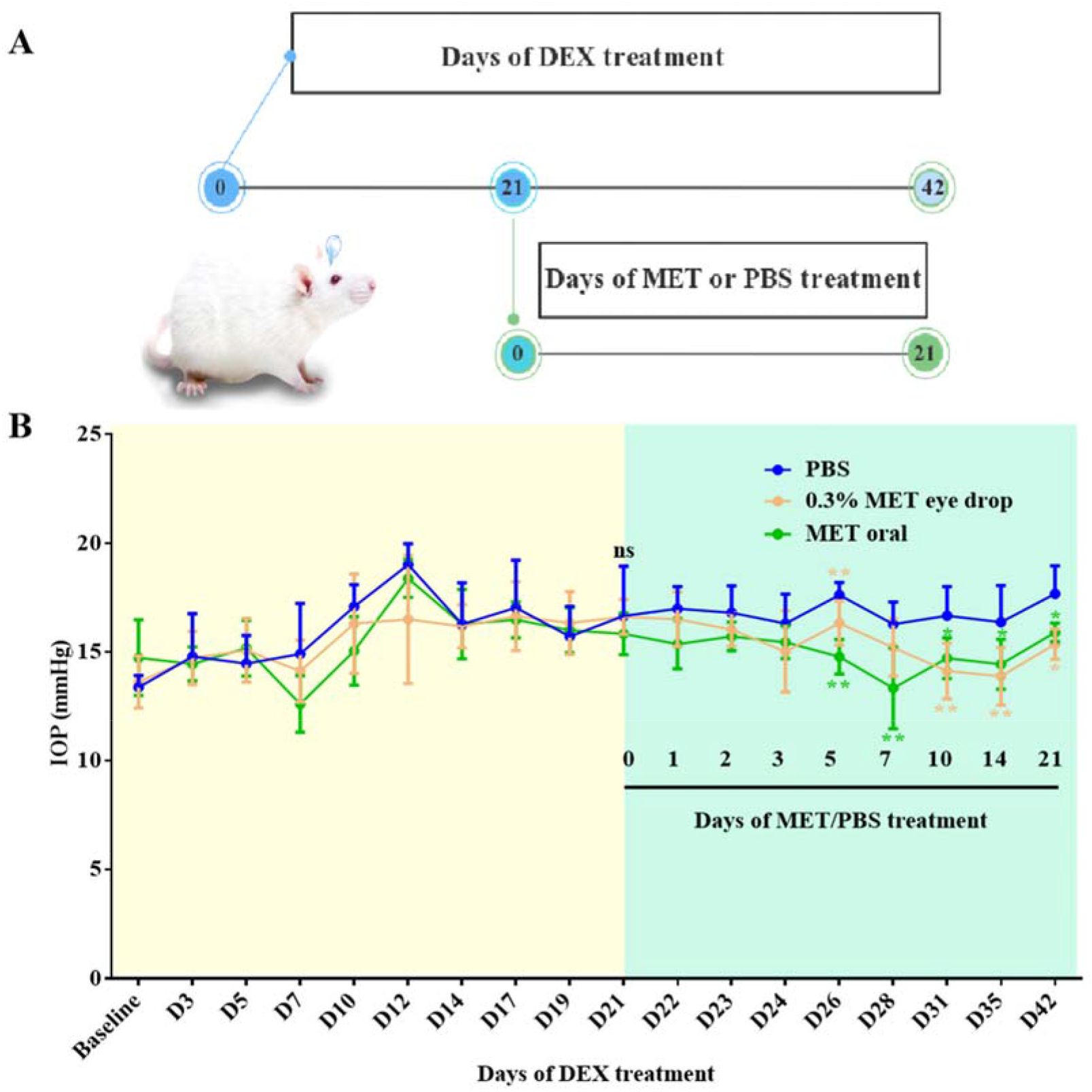
Effect of metformin (MET) on OHT mice model. A. Experimental process overview. B. MET effectively reversed the IOP in steroid-induced OHT mice models. \**p* < 0.05, \*\**p* < 0.01, ns: non-significance.

### Metformin attenuated the steroid-induced TM cytoskeleton damages in vivo

As shown in Figure 3, metformin, both eye drops and oral intake, improved fibrosis and the intensity of phalloidin labelling of F-actin in TM tissue in steroid-induced OHT C57BL/6 mice (Figure 3A-B). Quantitative comparison showed a significant difference in the number of α-SMA-, TGF-β-, and fibronectin (FN)-positive TM cells between MET-treated and control mice in a time-dependent manner (Figure 3C-J).

**Figure 3.**
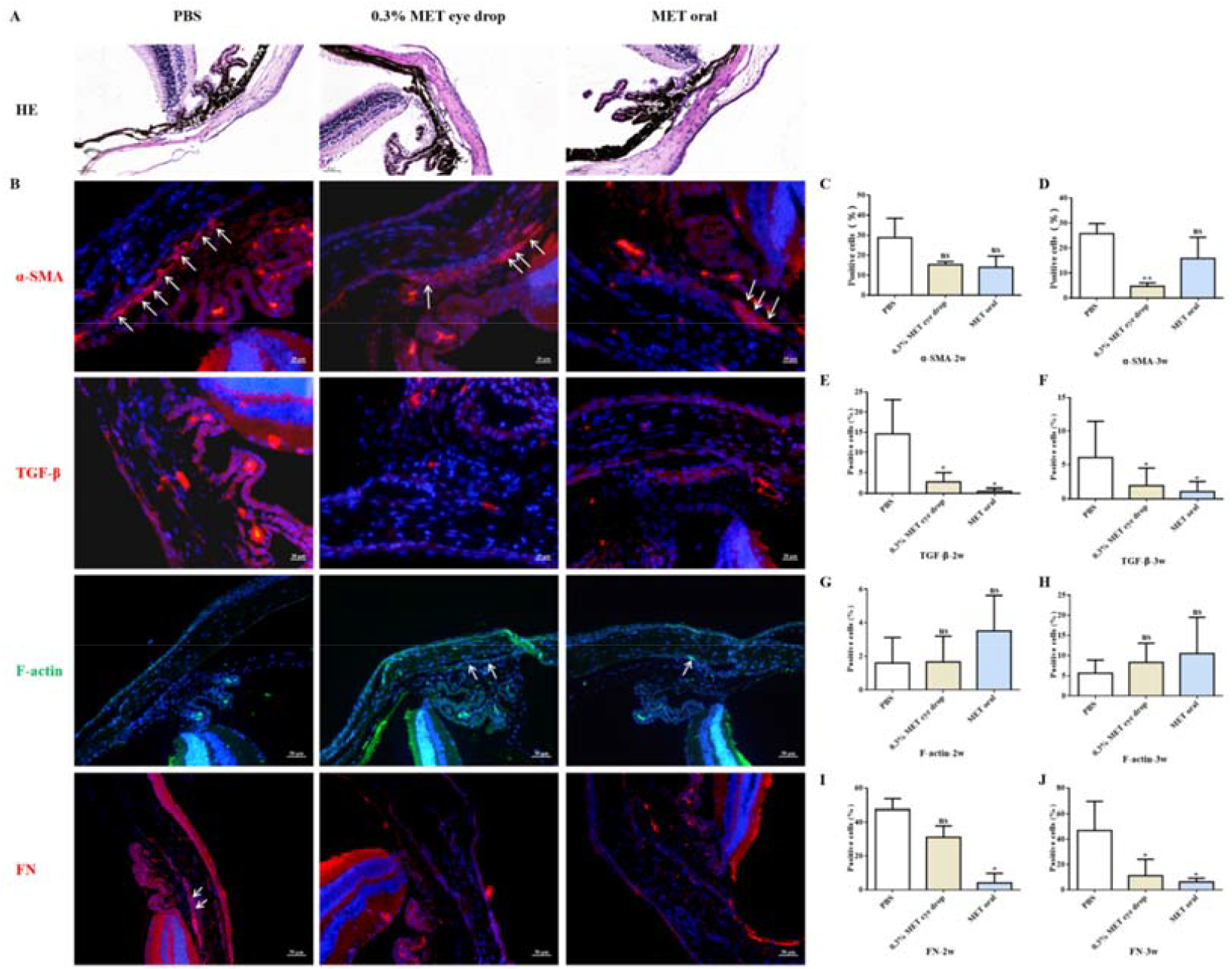
MET decreased the expression of fibrotic markers in steroid-induced trabecular meshwork stiffening in mice. A-B. Representative images of HE (A) and fibrotic markers (B). C-D. Quantification of α-SMA of the models after 2 weeks (C) and 3 weeks (D) of MET treatment. E-F. Quantification of TGF-β of the models after 2 weeks (E) and 3 weeks (F) of MET treatment. G-H. Quantification of F-actin of the models after 2 weeks (G) and 3 weeks (H) of MET treatment. I-J. Quantification of fibronectin (FN) of the models after 2 weeks (I) and 3 weeks (J) of MET treatment. \**p* < 0.05, \*\**p* < 0.01, ns: non-significance. White arrows indicate the representative positive cells.

Although not statistically significant, metformin treatment promoted the cytoskeleton recovery of steroid-induced TM cell damage, as confirmed by the upregulation of F-actin. We concluded that the IOP-lowering effect of metformin in this steroid OHT model can be largely explained by the attenuation of fibrotic alterations and rearrangement of the cell skeleton at sites of TM or trabecular outflow pathways.

### Protection effects of metformin on TM in vitro

To test the effect of metformin on TM *in vitro*, we performed studies on HTMC. WB showed that the expression of myocilin, a glucocorticoid-inducible gene in the HTMC, increased after DEX treatment (Figure 4B), confirming that the cells had characteristics of trabecular meshwork cells.

**Figure 4.**
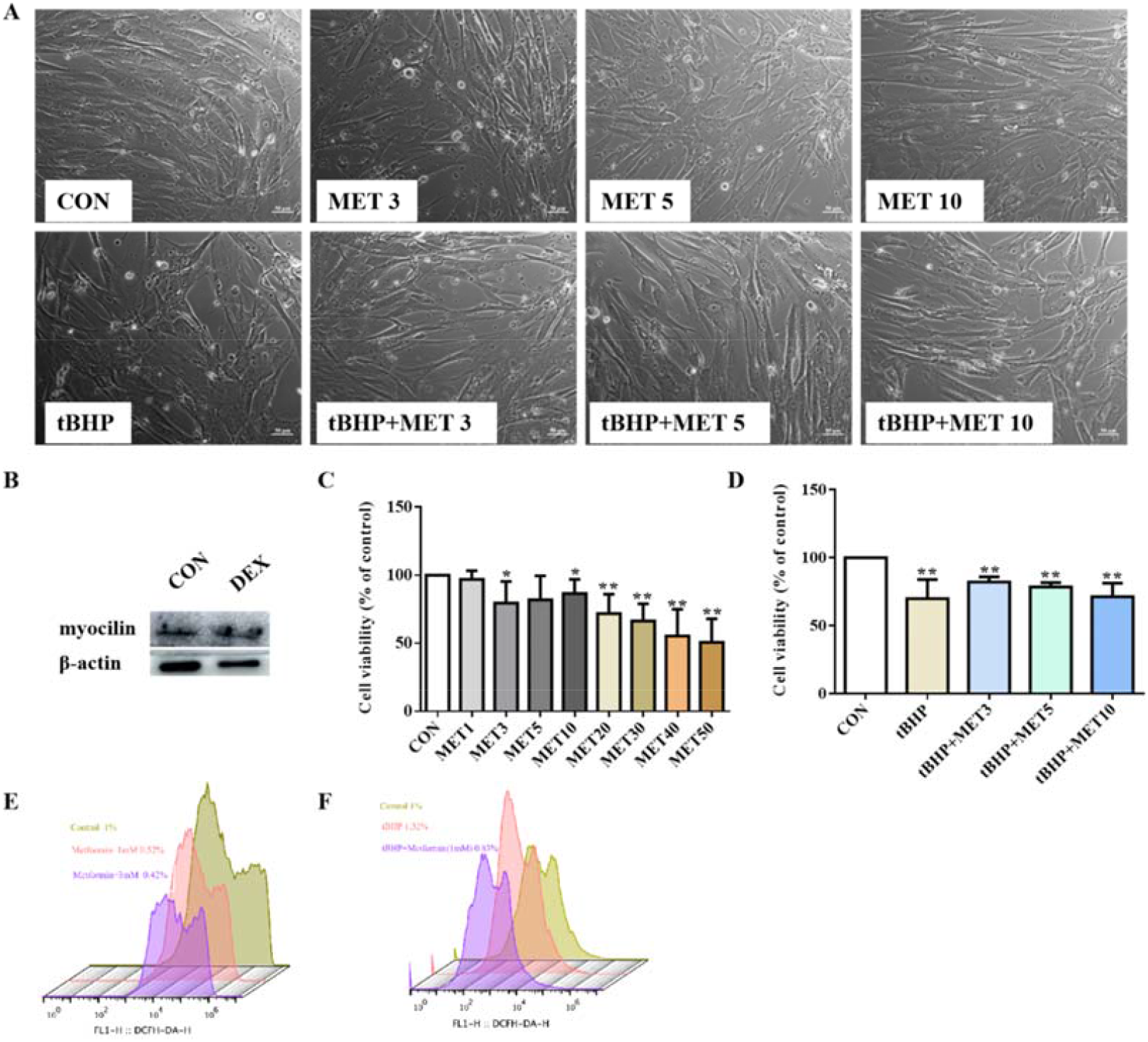
Low dose of metformin reversed the disarranged morphology of HTMC. A. HTMC were treated with metformin for 24 h with or without pre-treatment of tBHP for 1 h. Representative images of cell distribution and morphology photographed by inverted microscopy. B. The expression of myocilin after DEX treatment in HTMC. C-D. The relative HTMC viability after exposure to MET with different concentrations. Cell proliferation was measured using the CCK8 assay. E-F. The relative ROS levels were assayed via FCM and the results showed that metformin reduced the ROS production of HTMC induced by tBHP. \*\**p* < 0.01. \**p* < 0.05.

Higher doses (≥ 10 mM) of metformin decreased the number of HTMC, but this inhibitory effect was not evident at lower doses (< 10 mM) (Figure 4C). To verify the protective role of metformin in HTMC, cells were pre-treated with 100 μM tBHP for 1 h to induce chronic oxidative stress (*Tang, et al.,2013*), and subsequently exposed to low doses of metformin (3, 5, and 10 mM) (shortened as L-MET in this study). The results showed that L-MET significantly reversed the inhibitory effect induced by tBHP (Figure 4A, 4D).

Furthermore, cells treated with L-MET showed less intracellular ROS signals compared to the control, and 1 mM metformin reduced tBHP-induced ROS production. These results were confirmed by FCM analysis (Figure 4E-F). These results indicate that metformin reversed oxidative damage to HTMC.

### Metformin restored the tBHP induced cytoskeleton destruction in HTMC and activated integrin/ROCK signals

Actin filaments play key roles in cortical polarisation and asymmetric spindle localisation during phagocytosis in HTMC. F-actin was stained to determine whether actin dynamics were involved in tBHP-induced cellular dysfunction. As shown in Figure 5A, F-actin was evenly accumulated in the cytoplasm with a robust fluorescent signal in control HTMC. Exposure to L-MET did not change the morphology of HTMC as observed by inverted phase-contrast microscopy. Contrarily, HTMCs exposed to 1 h-tBHP displayed intermittent distribution of actin filaments with faded fluorescent signals. Additionally, tBHP treatment significantly destroyed HTMC, manifesting as various morphology-aberrant cells with misaligned cytoplasm. These pathological changes could be partially rescued by L-MET, implying that metformin could restore the dynamic instability from oxidative damage. Furthermore, WB results (Figure 5B-C) showed that L-MET significantly activated the integrin/ROCK pathway by upregulating integrin, ROCK, pAMPK, and F-actin, and it was more pronounced in the 3 mM and 5 mM doses.

**Figure 5.**
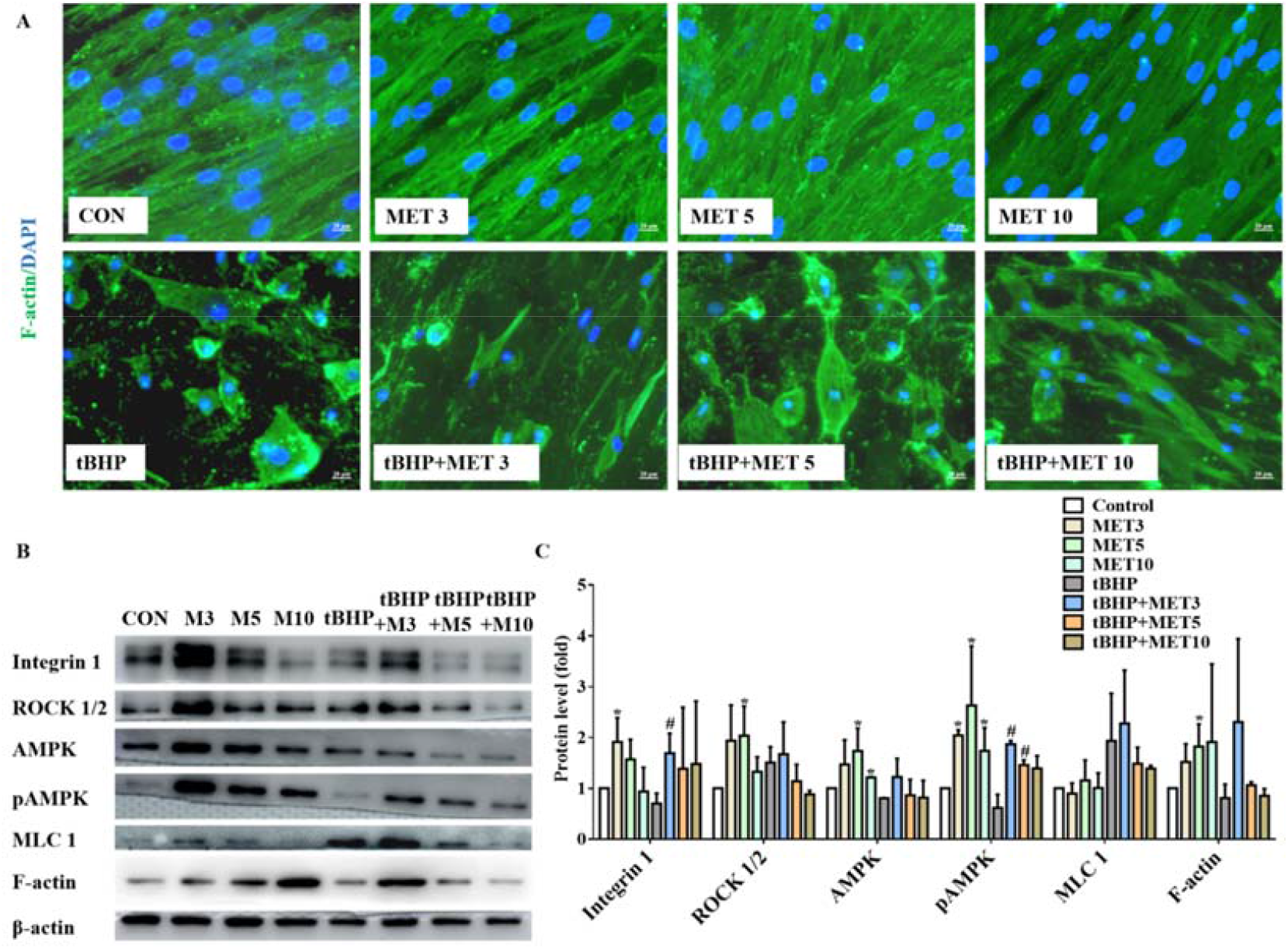
Metformin promoted the recovery of tBHP-induced cytoskeleton damages (A) and activated intergrin/ROCK pathway (B) in HTMC. C. The quantitative protein levels of B. * *p* < 0.05 (comparison with the control), # *p* < 0.05 (comparison with the tBHP treated group).

The ultrastructures of HTMC were examined by transmission electron microscope (TEM) (Figure 6). We observed a significant reduction in the amount and density of microfilaments in the tBHP treated cells and these changes were recovered after L-MET treatment.

**Figure 6.**
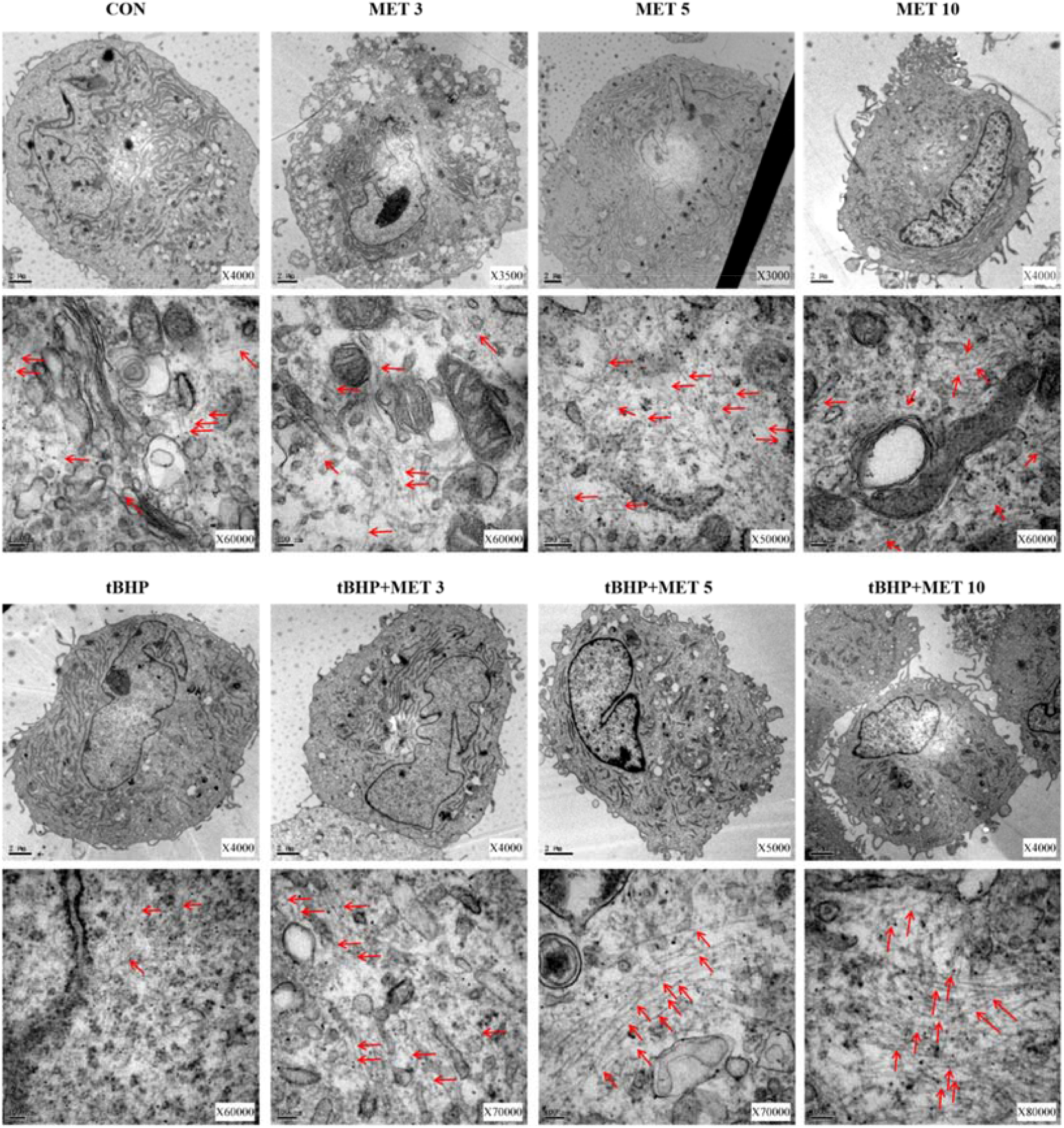
Metformin partially normalized the damaged microfilaments of HTMC induced by tBHP. Red arrows indicate the representative microfilaments imaged by TEM. induced by tBHP. Red arrows indicate the representative microfilaments imaged by TEM.

## Discussion

The major observations of the current study were the three significant phenotypic changes induced by metformin in OHT mouse eyes and HTMC. The first was the IOP lowering effect in steroid-induced OHT mouse eyes. The second was a reversal of the skeletal destruction of TM cells, both *in vivo* and *in vitro*. Third, there was a significant decrease in the accumulation of fibrotic markers, namely α-SMA, TGF-β, and fibronectin, in TM tissues. These results suggest a protective effect of metformin in TM, probably via promoting cytoskeleton recovery through the integrin/ROCK pathway.

Steroid-induced OHT in mice was generated (*Wang, et al.,2018*) with the characteristics of TM stiffening and elevated IOP (*Li, et al.,2019*). In line with previous studies (*Clark, et al.,2001*; *Johnson, et al.,1997*; *Johnson, et al.,1990*), our model suggested that the stiffness of AHO closely matched that of OAG in humans, including the deposition of ECM in the TM, disordered cytoskeleton, increased AHO resistance, and elevated IOP. Similar to the study by Zode et al.,*(Zode, et al.,2014)* our topical DEX elevated the IOP by 0.3–6 mmHg after 3 weeks of induction.

TM tissues play an important role in the regulation of AHO function and IOP (*Wang, et al.,2018*). TM cells have the properties of extracellular debris phagocytosis and ECM degradation. If these functions are disturbed, the balance of TM in keeping AHO free of obstructive debris is disrupted (*Last, et al.,2011*). It is noteworthy that metformin appeared to restore TM biomechanical properties in our steroid-induced OHT model after 5 days of treatment, and this restoration was accompanied by significant downregulation of ECM proteins in TM.

Moreover, as evidenced by IF and WB analyses, L-MET rearranged the disordered cytoskeleton of HTMC. This finding was supported by a study conducted by Li et al (*Li, et al.,2020*), who showed that 50 μM metformin protected oocytes against cytoskeleton destruction. Interestingly, the time course of structural changes in the TM was consistent with the observed pharmacodynamics of metformin on IOP, significantly decreasing 2 and 3 weeks of OHT because of MET treatment, which was consistent with the known effects of other ROCK regulators (*Lin, et al.,2018*; *McMurtry, et al.,2010*). Taken together, our results indicate that L-MET promotes the regeneration of abnormal HTMC and enables the recovery of TM function; this should be confirmed in future studies.

As confirmed by WB analysis, complementation with L-MET treatment upregulated integrin, pAMPK, ROCK, and F-actin signals. These *in vitro* findings indicate that metformin might elicit its protective effects through the regulation of the integrin/ROCK pathway in HTMC. The *in vivo* observations in the OHT model again support the importance of the integrin/ROCK pathway in TM tissue protection. Generally, these signals mediate cellular biomechanical tension through actomyosin cytoskeletal tension, ECM synthesis, assembly, and degradation (*Nakajima, et al.,2005*; *Pattabiraman and Rao,2010*; *Rao, et al.,2001*).

Inconsistent with the previous studies indicating that the AMPK activator suppressed ROCK activity, our study revealed a novel mechanism of metformin action within which activation of both AMPK and ROCK signals sequentially upregulated F-actin expression in TM. We hypothesised that ROCK regulators might protect TM via different mechanisms, either by inhibiting ROCK signals when increased AHO resistance is attributed to TM stiffness or by activating the ROCK pathway to promote cytoskeleton formation when it is damaged. In the current study, we presumed that metformin lowered IOP through short-term changes in TM cell morphology and adhesion by regulating integrin/ROCK signalling. In addition, recovery to normal IOP prevents further TM cell loss, whereas metformin promotes the maintenance of cytoskeleton morphology and repair after injury; therefore, IOP is maintained at a normal level for an extended period.

Generally, these data emphasise the protective properties of metformin in the conventional outflow pathway, consistent with a large body of literature on other tissues (*Rangarajan, et al.,2018*; *Yi, et al.,2021*; *Zhao, et al.,2021*). In addition to metformin-mediated changes in ECM turnover, the observed changes in ECM composition and amount may also be due to the metformin-mediated opening of flow pathways and the consequential removal of ECM. Additionally, the observed reduction in ROS content in HTMC treated with L-MET was in line with earlier studies displaying its protective effect in several kinds of cells (*Ghasemnejad-Berenji, et al.,2018*; *Huang, et al.,2015*; *Louden, et al.,2014*). In any case, therapy for steroid-induced OHT, which inhibits the cycle of damage and promotes cell remodelling by restoring function to a diseased tissue, offers a potential benefit for patients.

In summary, promotion of TM cytoskeleton recovery by metformin treatment may be an underlying mechanism for IOP reduction and AH outflow increase. A thorough investigation of the mechanisms by which cytoskeletal recovery is promoted will improve our understanding of the therapeutic mechanism of metformin. If metformin is confirmed to promote damaged cytoskeleton recovery and maintenance in the human body, it may have clear therapeutic effects on the main sites of glaucoma pathogenesis, which will broaden their application in the field of glaucoma. In conclusion, we revealed that integrin/ROCK activation reprograms metabolism in TM cells by enhancing the cytoskeleton and downregulating excessive ECM proteins.

## Materials and methods

| Reagent type<br>(species) or<br>resource | Designation | Source or<br>reference | Identifiers | Additional<br>information |
| --- | --- | --- | --- | --- |
| Strain,<br>strain background<br>(Mus musculus) | C57BL/6J | Zhejiang Vital River Experimental<br>Animal Technology Co. LTD<br>(Charles River lab China) |  |  |
| Cell culture | HTMC | BNCC(Shanghai, China) | 338506 |  |
| Commercial assay<br>or kit | CCK8 kit | APE BIO |  |  |
| Commercial assay<br>or kit | DCFH-DA probe | Beyotime Biotechnology, Shanghai,<br>China |  |  |
| Drug | Dexamethasone | Shanghai Macklin Biochemical Co.,<br>Ltd, China |  | Cell:500 nM;<br>Animal:0.1% |
| Drug | Metformin | Sigma-Aldrich, St. Louis, USA |  | Cell:0, 1, 3, 5,<br>10, 20, 30, 40,<br>and 50 mM;<br>Animal:0.3%,<br>7.5 µg/day<br>per mouse |
| Drug | tBHP | Damas-beta, China |  | Cell:100 µM |
| Antibody | Anti-myocilin((rabbit monoclonal) | Abcam | 41552 | WB(1:1000) |
| Antibody | Anti-integrin 1(rabbit monoclonal ) | Abcam | Ab179471 | WB(1:1000) |
| Antibody | Anti-AMPK(rabbit monoclonal ) | CST | 2795T | WB(1:1000) |
| Antibody | Anti-pAMPK(rabbit monoclonal) | CST | 2535T | WB(1:1000) |
| Antibody | Anti-ROCK1/2(rabbit monoclonal) | Abcam | 45171 | WB(1:1000) |
| Antibody | Anti-MLC1(rabbit monoclonal ) | Abcam | Ab186436 | WB(1:1000) |
| Antibody | Anti-F-actin(mouse monoclonal ) | Abcam | Ab130935 | WB(1:1000)<br>IF (1:500) |
| Antibody | Anti-β-Actin(mouse monoclonal) | CST | 3700S | WB(1:1000) |
| Antibody | Anti-α-SMA(rabbit monoclonal) | CST | 19245 | IF(1:500) |
| Antibody | Anti-TGF-β(rabbit monoclonal) | Abcam | 215715 | IF(1:500) |
| Antibody | Anti-fibronectin(rabbit monoclonal) | Abcam | Ab268020 | IF(1:500) |
| Antibody | Anti-Rabbit IgG H&L,F(ab')<sub>2</sub> Fragment(Alexa Fluor 594 Conjugate) | CST | 8889S | IF(1:500) |
| Antibody | Anti-Mouse IgG H&L,F(ab')<sub>2</sub> Fragment(Alexa Fluor 488 Conjugate) | CST | 4408S | IF(1:500) |
| Antibody | Anti-mouse IgG, | CST | 7076S | WB(1:10000) |
|  | HRP-linked Antibody |  |  |  |
| Antibody | Anti-rabbit IgG,<br>HRP-linked Antibody | CST | 7074S | WB(1:10000) |
| Other | DAPI | Beyotime Institute of Biotechnology,<br>Shanghai, China |  | IF(1:500) |
| Software,<br>algorithm | SPSS software | SPSS software | Version<br>22.0 |  |
| Software,<br>algorithm | GraphPad Prism | GraphPad Software | Version 7.0 |  |
| Software,<br>algorithm | ImageJ | National Institutes of Health, USA | Version<br>1.52a |  |

### Animals

All the animals were treated in accordance with the principles of the Declaration of Helsinki and in compliance with the Association for Research in Vision and Ophthalmology (ARVO) Statement. All experiments were approved by the Institutional Animal Care and Use Committee of the Wenzhou Medical University (wydw2022-0209). Healthy C57BL/6J mice (age: 6–8 weeks, male) were used in this study. The mice were purchased from Zhejiang Vital River Experimental Animal Technology Co. LTD (Charles River lab China), bred/housed in clear cages, and kept in housing rooms at 21°C with a 12 h:12 h light: dark cycle.

### OHT animal model and drug treatments

The animal experiments were performed in two consecutive steps. In the first step, OHT C57BL/6J mouse model was induced by topical 0.1% dexamethasone phosphate (DEX) twice daily (8–9 AM and 5–6 PM) as described previously (*Li, et al.,2021*) and sterile phosphate buffer saline (PBS) was used as a vehicle control. The second step was conducted 3 weeks after steroid induction. The successfully created OHT mice models with elevated IOP were randomly divided into three groups: control (PBS eye drops, twice daily), 0.3% MET eye drops (twice daily), and MET oral groups (7.5 μg/day for each mouse). In this step, the mice received an additional 3 weeks of supplemented drug delivery.

### IOP measurements

Briefly, mice were anaesthetised using gaseous isoflurane (approximately 2 min) and topical alcaine (Alcon-couvreur n. v, Rijksweg, Puurs Belgium). IOP was measured using a rebound tonometer (icare TONOVET; Vantaa, Finland).Each recorded IOP was the average of five measurements and three IOP readings were recorded for the same eye to calculate the mean value. IOP measurements were conducted every 2–3 days (Monday, Wednesday, and Friday, between 2 PM and 3 PM).

### Weight Measurement

The anaesthetised mice were gently placed on digital electronic scales (Electronic Scale, Kunshan, China) to measure their weight. The effective reading of body weight (BW) was recorded to an accuracy of 0.01 g. The BW of each mouse was calculated from the average of the three test values.

### Cell culture and treatment

Human TM cells (HTMC) were purchased from BNCC (338506, Shanghai, China) and cultured in DMEM/F12 medium (Cytiva, HyClone Laboratories, Logan, Utah,) containing 10% foetal bovine serum (FBS, BI, USA) and antibiotics (100 U/mL penicillin and 100 μg/mL streptomycin, Gibco, Life Technologies Corporation, NY, USA) at 37°C and 5% CO2. To identify the characteristics of HTMC, cells were treated with 500 nM dexamethasone (DEX) (Shanghai Macklin Biochemical Co., Ltd, China) for 7 days, and the expression of myocilin was evaluated by western blotting (WB) (*He, et al.,2019*). For drug testing, tBHP (Damas-beta, China) and metformin (Sigma-Aldrich, St. Louis, USA) were dissolved in DMEM/F12. HTMC were pre-treated with tBHP solution for 1 h to induce oxidative damage, followed by 24-h incubation in normal culture medium containing metformin at certain concentrations.

### Cell viability

Cell viability was measured using the CCK8 assay kit (APE BIO), following the manufacturer’s instructions. The cells were seeded in 96-well plates and exposed to increasing concentrations of metformin (0, 1, 3, 5, 10, 20, 30, 40, and 50 mM) for 24 h. The cell viability was determined by measuring the optical density at 490 nm using an absorbance microplate reader (SpectraMax 190; version 7.1.0, Molecular Devices, California, USA).

### Detection of intracellular reactive oxygen species levels

Intracellular ROS levels were determined using a 2′,7′-dichlorofluorescein diacetate (DCFH-DA) probe (Beyotime Biotechnology, Shanghai, China) according to the manufacturer’s instructions and a previous study (*Xu, et al.,2020*). The values were normalised to signals from the control group.

### Western blotting

Cells were lysed 24 h after drug treatment as previously described (*Xu, et al.,2020*). After gel separation and membrane transfer, myocilin (Abcam), integrin 1 (Abcam), AMPK (CST), pAMPK (CST), ROCK1/2 (Abcam), MLC1 (Abcam), and F-actin (Abcam) were detected. β-Actin (CST) was used as the loading control. Western blot membranes were developed using a chemiluminescence detection system (Amersham Imager 680RGE; GE Healthcare Bio-Sciences AB, Sweden, Japan).

### Histology, immunostaining and transmission electron microscope (TEM)

At the time of harvest, the mice were re-anesthetized, and the eyes were fixed in 4% paraformaldehyde overnight, embedded in paraffin or optimum cutting temperature compound in sagittal axis. The sections were incubated overnight with primary antibodies at 4°C according to the manufacturers’ instructions (*Xu, et al.,2020*).α-SMA (CST), TGF-β (Abcam), F-actin (Abcam), fibronectin(Abcam)were detected. The secondary antibodies were goat anti-rabbit or anti-mouse (CST) at a 1:500 dilution. Sections were subsequently incubated with 2-(4-Amidinophenyl)-6-indolecarbamidine dihydrochloride (DAPI, Beyotime Institute of Biotechnology, Shanghai, China) for 5 min to stain the nuclei, washed, and then mounted.

The cells were fixed with 4% paraformaldehyde for 15 min, permeabilised with 0.25% Triton X-100 for 20 min, and blocked with 5% bovine serum albumin for 1 h at room temperature. After blocking, they were incubated overnight with primary antibodies at 4°C. The next day, the cells were rinsed and incubated with secondary antibodies conjugated to Alexa Fluor (CST) for 1 h at room temperature. Fluorescent images were obtained using a confocal microscope (X-Cite Series 120, Lumen Dynamics Group Inc., Canada).

For electron microscopy studies, cells was fixed in 2.5% glutaraldehyde (Shanghai Macklin Biochemical Co., Ltd, China) and embedded in Epon resin and 80 nm sagittal thin sections were cut through iridocorneal tissues using an ultramicrotome (Power Tome-XL, RMC Products, USA). Sections were stained with uranylacetate/lead citrate and examined with a transmission electron microscope (HITACHI, H-7500).

### Image analysis

Immunohistochemical images were obtained using a microscope (ECLIPSE 80i, Nikon) and analysed using the NIS-Elements Imaging Software (3.22.00; Build 700, LO, USA). For quantification, high-power fields (400× magnification) of the AHO from each model were captured. ImageJ (v1.52a, National Institutes of Health, USA) was used to quantify the positively stained cells.

### Statistical analysis

Data analysis was conducted using SPSS (version 22.0) software and GraphPad Prism (version 7.0). Cell-based and animal-based experiments included at least three biological replicates. Continuous data are summarised as mean±standard deviation.

The Student’s t-test and ANOVA were used to test the differences in continuous variables. A paired t-test was used to compare IOP changes from the baseline. Statistical significance was set at *p* value < 0.05.

## Data availability

All data generated or analysed during this study are included in the manuscript and supporting files. Source data files have been provided for Figures 1, 2, 3, 4 and 5.

